# Genetically Encodable *in situ* Gelation Redox-Responsive Collagen-Like Protein Hydrogel for Accelerating Diabetic Wound Healing

**DOI:** 10.1101/2023.05.29.542680

**Authors:** Shuang Jia, Jie Wang, Xiaojie Wang, Xing Liu, Shubin Li, Yimiao Li, Jiaqi Li, Jieqi Wang, Shad Man, Zhao Guo, Yinan Sun, Zhenzhen Jia, Liyao Wang, Xinyu Li

## Abstract

Genetically encoded collagen-like protein-based hydrogels have demonstrated remarkable efficacy in promoting the healing process in diabetic patients. However, the current methods for preparing these hydrogels pose significant challenges due to harsh reaction conditions and the reliance on chemical crosslinkers. In this study, we present a genetically encoded approach that allows for the creation of protein hydrogels without the need for chemical additives. Our design involves the genetic encoding of paired-cysteine residues at the C- and N-terminals of a meticulously engineered collagen-like recombination protein. The protein-based hydrogel undergoes a gel-sol transition in response to redox stimulation, achieving a sol-gel transition. We provide evidence that the co-incubation of the protein hydrogel with 3T3 cells not only enhances cell viability but also promotes cell migration. Moreover, the application of the protein hydrogel significantly accelerates the healing of diabetic wounds by upregulating the expression of COL-1α and CK-14, while simultaneously reducing oxidant stress in the wound microenvironment. Our study highlights a straightforward and chemical-free strategy for the preparation of redox-responsive protein hydrogels. Importantly, our findings underscore the potential of this hydrogel system for effectively treating diabetic wounds, offering a promising avenue for future therapeutic applications.

## 1. Introduction

Diabetes-induced hyperglycemia leads to the chronic wounds characterized by bacterial infections[1, 2], oxidant stress[3, 4], inflammatory reactions[5, 6] and low angiogenesis[7, 8]. Effective therapy for diabetic wounds (DW) is crucial to improve the quality of life for diabetic patients, as the associated costs of wound care, including frequent dressing changes and hospitalizations, impose a significant economic burden on both patients and healthcare systems[3, 6, 9].

Protein-based hydrogels have emerged as promising materials for diabetic wound therapy due to their biocompatibility, bioactivity, and resemblance to natural skin tissues[8, 10]. In recent years, both natural proteins, such as elastin[11, 12], silk[10], collagen[13], have been utilized to develop hydrogels for diabetic wound healing. To further enhance the therapeutic efficacy, growth factors[12], cytokines[5] or micro-RNAs[14] are commonly incorporated into the hydrogel system. However, it is important to note that these bioactive additions are prone to degradation and metamorphosis, necessitating the use of mild crosslinking approaches during hydrogel preparation.

While various crosslinking methods, including photo-crosslinking, thermal-induced crosslinking, and chemical crosslinkers, have been developed for protein-based hydrogels, they still pose challenges due to the sensitivity of proteins to factors such as pH, temperature, or ionic concentration[15]. Some of these methods can adversely affect the structural and functional properties of the proteins. For example, high-temperature and chemical coupling strategies used for the preparation of BSA-based hydrogels result in the unfolding of BSA and loss of its functions [16, 17]. Additionally, certain crosslinking methods require the modification of functional groups, which can alter the structure and properties of the proteins. Although supramolecular crosslinker methods have been developed for protein crosslinking [18, 19], they often result in weaker molecular networks and unstable response activities. Therefore, there is a need to develop crosslinking methods that allow for spontaneous reactions under mild conditions, while maintaining stimulated response activities, for the preparation of protein-based hydrogels.

The remarkable ability of cysteine, an amino acid containing a sulfhydryl group, to form disulfide bonds under oxidizing conditions [20], has inspired researchers to develop protein-based hydrogels crosslinked through disulfide bonds [15, 20]. These pioneering studies have highlighted the potential of cysteine-based crosslinking approaches for *in situ* protein hydrogel preparation. The utilization of cysteine-based crosslinking provides a versatile strategy for constructing protein-based hydrogels and opens up new opportunities for practical applications. Importantly, the ability to engineer specific sequences with cysteine residues enables precise control over hydrogel properties, including mechanical strength and degradation rates [8, 15, 20]. The innovative research conducted by these pioneering groups underscores the significance of cysteine in the development of protein-based hydrogels and further establishes it as a valuable tool in the field. This knowledge contributes to the ongoing advancement and exploration of novel protein-based hydrogel systems.

Genetically encoded collagen-like proteins (CLPs) have been engineered and utilized for the fabrication of hydrogels with exceptional wound healing properties [21-23]. Various crosslinking methods, such as EDC-NHS[24], supramolecular self-assembly[19], and non-covalent crosslinked with polysaccharides[25], have been reported for the preparation of CLP-based hydrogels. However, the development of a recombination CLP that can undergo *in situ* gelation without the need for chemical additives remains a challenge. Given their high productivity, “blank collagen template” characteristics, and suitability as building blocks for protein-based hydrogel synthesis, exploring spontaneous crosslinking approaches under mild conditions holds great promise for advancing CLP-hydrogel development.

In this study, we introduce two meticulously designed collagen-like proteins derived from Scl-2, serving as the foundation for a genetically encoded protein-based hydrogel: C1-eCLP3-C1 and C2-eCLP3-C2. These hydrogels demonstrate the unique ability to undergo spontaneous gelation under oxidizing conditions and undergo gel-sol transition in reducing environments. Importantly, our findings highlight the significant potential of these hydrogels in accelerating the healing process of diabetic wounds. Thus, these hydrogels hold promise for diverse applications in regenerative medicine and tissue engineering. Furthermore, the crosslinking strategy employed in this study provides a valuable platform for the development of redox-responsive protein hydrogels and the design of advanced hydrogel systems.

## 2. Results

### 2.1 Protein Construct Design

The protein sequences utilized in this study were designed based on the Scl-2 protein from *Streptococcus pyogenes*[21, 26]. To construct the protein, the native V-domain, a globular domain, was fused to the N-terminal of three repeated collagen-like domains **(Fig1. a)**. To enhance the stability of the protein, the sequences of the collagen-like domains were modified to contain a higher proline content, and two stabilizing sequences, KGD and KGE, were incorporated[21, 27]. Furthermore, two CPPC peptides were introduced at the N- and C-terminal of the collagen-like domains to further reinforce the stability, as these peptides have been demonstrated to stabilize the triple helix structure of collagen-like domains[25] **(Fig1. a)**. Additionally, RGD sequences were integrated into the protein to facilitate cell adhesion **(Fig. 1a)**. To enable *in situ* gelation of the protein, one or two paired cysteine residues with glycine (CGG) were introduced at the N- and C-terminal of the protein, named C1-eCLP3-C1 and C2-eCLP3-C2, respectively. The paired cysteines could provide stable crosslinking sites for hydrogel formation **(Fig1. c)**. The protein sequence was cloned into a pET-28a vector and expressed in BL21(DE3) *E. coli* cells. Following purification and concentration, the hydrogel was formed within 48 hours at 4°C or upon treatment with 200 μM H_2_O_2_ **(Fig1. b)**. Under oxidizing conditions, the thiol groups of the designed paired cysteines crosslink with one another, facilitating gel formation **(Fig. 1c)**.

**Figure 1.**
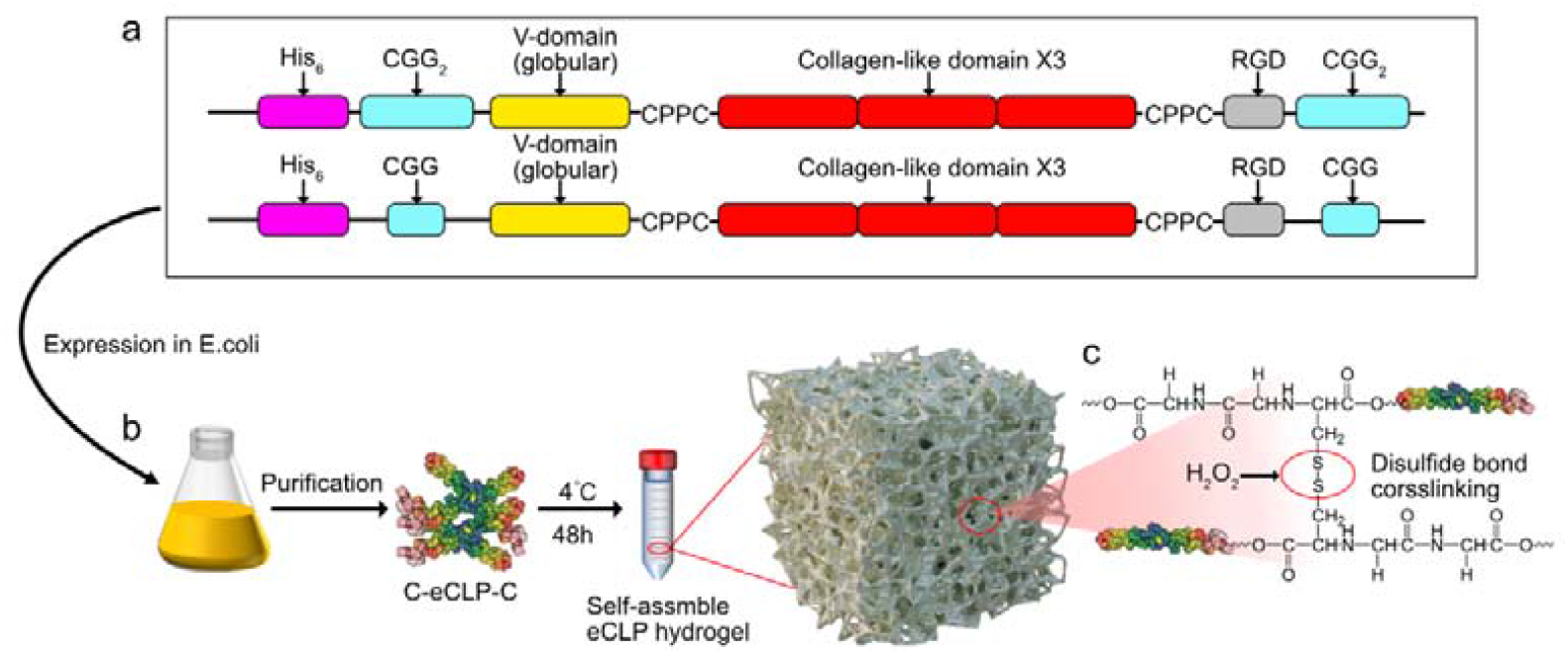
Protein design and hydrogel formation. a. Schematic representation of the protein construction. b. Expression of the proteins and formation of the hydrogel. c. Illustration of the crosslinking mechanism in the hydrogel.

### 2.2 Synthesis and Molecular Characterization of Proteins

The protein sequences were subjected to structural prediction using Alphafold **(Fig. 2a)**. The results indicated that the paired cysteines are located on the outer surface of the protein structures and have sufficient spatial distance from each other **(Fig. 2a)**. This spatial arrangement suggests that they may offer suitable distances for crosslinking with other proteins. The proteins were then expressed using the BL21(DE3) *E.coli* expression system. After purification using Ni-NTA Agarose and HiTrap Capto Q columns. The purified proteins exhibited an apparent molecular weight ranging from 35 kDa to 48 kDa by SDS-page **(Fig. 2b)**. To further validate the identity of the purified protein, a Western blot analysis was performed using the anti-Histag primary antibody and the results confirmed the presence of the desired protein **(Fig. 2c)**. In addition, protein mass spectrometry was conducted to confirm the molecular weight of the protein, and the results indicated a molecular weight of approximately 36.4 kDa, which aligned with the expected size **(Supplementary Table.1)**. The mass spectrometry analysis also involved the digestion of the protein, resulting in the generation of 20 peptides. This extensive analysis enabled a comprehensive characterization of the protein’s sequence and structure **(Supplementary Table.1)**. The circular dichroism (CD) spectroscopy was employed to investigate the secondary structure of the proteins in different buffers. The CD results demonstrated that the secondary structure of the proteins remained stable in pH 7 and pH 8 citric acid/Na_2_HPO_4_ buffer as well as in PBS of pH 7 **(Fig.2.d)**. However, noticeable changes in the secondary structure were observed at pH 4 **(Fig.2.d)**.

**Figure 2.**
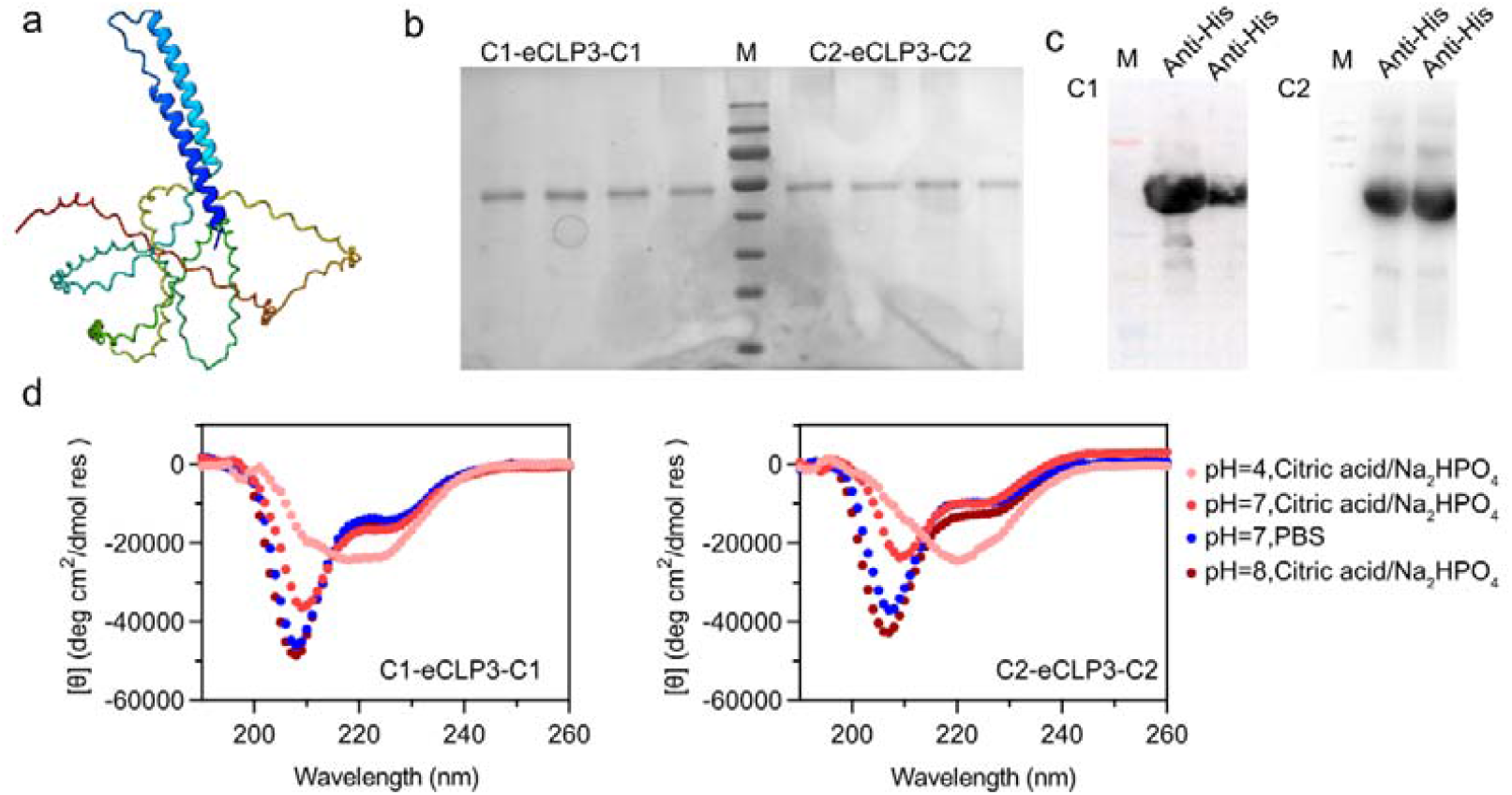
Molecular characterization of proteins. a. Alphafold-predicted protein structure of C2-eCLP3-C2. b. SDS-PAGE analysis of purified C1-eCLP3-C1 and C2-eCLP3-C2 proteins. c. Western blot images showing the presence of C1-eCLP3-C1 and C2-eCLP3-C2 proteins using anti-Histag antibody. d. Circular dichroism (CD) spectra of C1-eCLP3-C1 and C2-eCLP3-C2 proteins in different buffers.

### 2.3 Characterization of The Protein-based Hydrogel

The purified proteins exhibited the ability to form hydrogels after a 48-hour incubation at 4°C or upon the addition of 200 μM H_2_O_2_ followed by a 6-hour incubation **(Fig. 3a)**. These hydrogels displayed a soft jelly-like consistency and could be injected into desired areas. To further explore their redox-responsive characteristics, a final concentration of 200 μM DTT was introduced to the hydrogel, resulting in its transformation into a protein solution within 10 minutes **(Fig. 3b)**. Subsequently, upon the addition of 200 μM H_2_O_2_ to the protein solution, the solution gradually converted into a hydrogel over a period of 6 hours **(Fig. 3b)**. The hydrogels were subjected to lyophilization, and their chemical bonds were examined using FTIR, while their internal structure was analyzed using scanning electron microscopy (SEM) **(Fig. 3c, d)**. SEM revealed the presence of uniform porous structures, with the C2-eCLP3-C2 hydrogel displaying a more rigid structure compared to the C1-eCLP3-C1 hydrogel. This difference can be attributed to the increased number of cysteine residues, which provide additional crosslinking sites. Rheological analysis demonstrated that the storage modulus (G’) of the hydrogels consistently exceeded the loss modulus (G”), confirming their gel-like behavior and injectability **(Fig. 3e)**.

**Figure 3.**
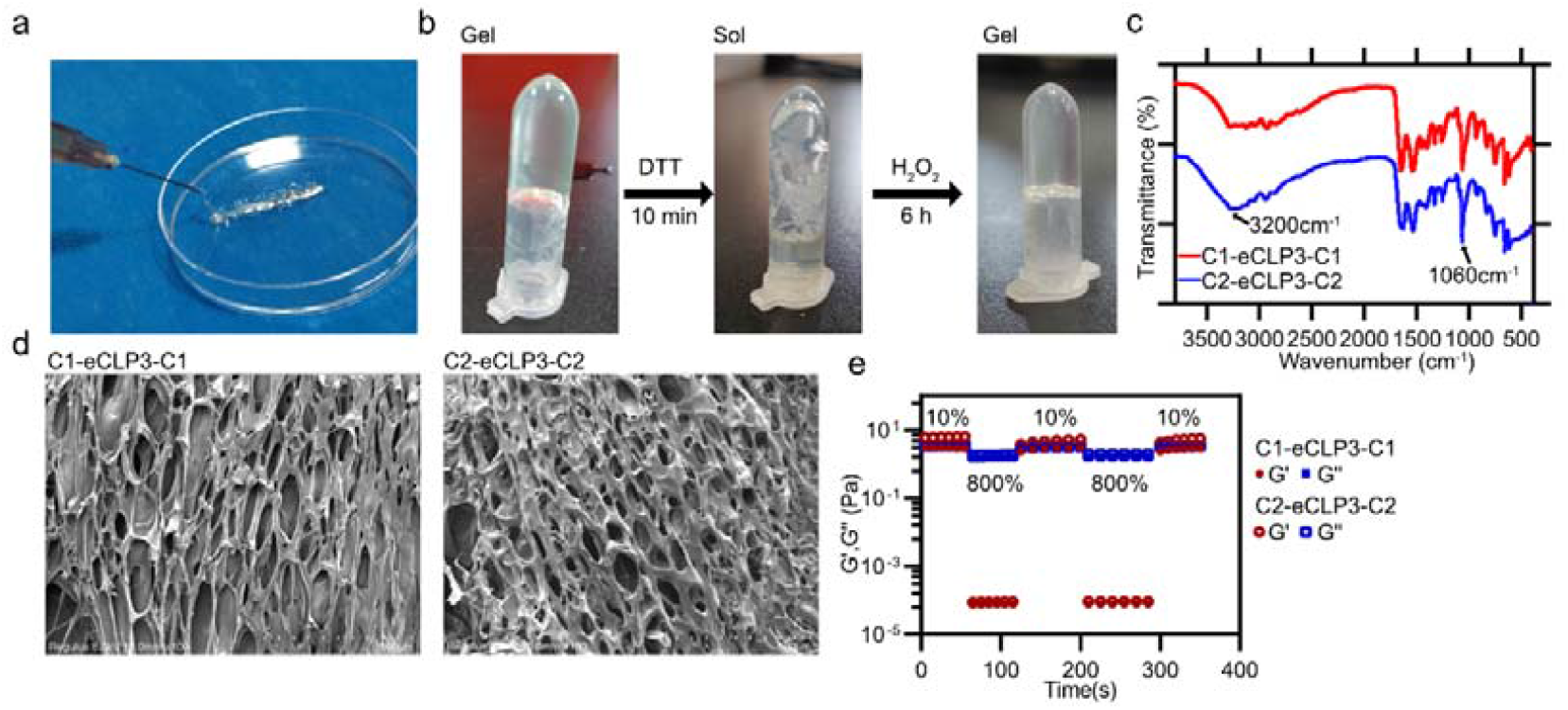
Morphology and characterization of the hydrogels. a. Photograph showing the injectable hydrogel. b. Gel-sol transition of the hydrogel in response to redox stimuli. c. FTIR spectra of the hydrogels. d. SEM images revealing the internal structure of the hydrogels. e. Dynamic step-strain measurements of the hydrogels at 37 °C, subjected to repeated deformation of 10% strain for 60 s and 800% strain for 60 s.

### 2.4 Biocompatibility Assessment

For clinical wound dressing applications, favorable biocompatibility is a crucial requirement for hydrogels[28]. In this study, we evaluated the biocompatibility of the hydrogels in vitro using live/dead staining **(Fig.4.a)**, the Cell Counting Kit (CCK-8) **(Fig.4.b)**, and a hemolysis test **(Fig.4.c)**. NIH-3T3 cells were co-incubated with hydrogels and native collagen at the same concentration as the positive control groups. After co-incubation, minimal dead cells were observed, and a higher number of live cells were found in the C1-eCLP3-C1 and C2-eCLP3-C2 hydrogel-treated groups compared to the native collagen and non-treated groups, indicating excellent cell viability **(Fig.4.a)**. The CCK-8 assay further confirmed the positive effect of hydrogel treatment on cell viability, with significant increases observed in the 1% and 10% treatment groups for C1-eCLP3-C1, C2-eCLP3-C2 hydrogels, and collagen, compared to the blank control group **(Fig.4.b)**. However, in the 30% treatment group, although cell viability was significantly increased, it was lower than that in the 10% treatment group **(Fig.4.b)**. This difference may be attributed to the high protein concentration in the 30% hydrogel, which could alter the osmotic pressure of the cell medium[29]. To assess the hemocompatibility of the hydrogels, a hemolysis assay was conducted[28, 30]. The C1-eCLP3-C1 and C2-eCLP3-C2 hydrogels showed no apparent hemolysis when incubated with fresh blood, indicating their non-hemolytic nature **(Fig. 4c)**. This finding is crucial as wound dressings inevitably come in contact with blood during clinical use[28]. Taken together, these results demonstrate that the C1-eCLP3-C1 and C2-eCLP3-C2 hydrogels exhibit favorable biocompatibility, support cell adhesion and growth, and are non-hemolytic, highlighting their potential for clinical applications as wound dressings.

**Figure 4.**
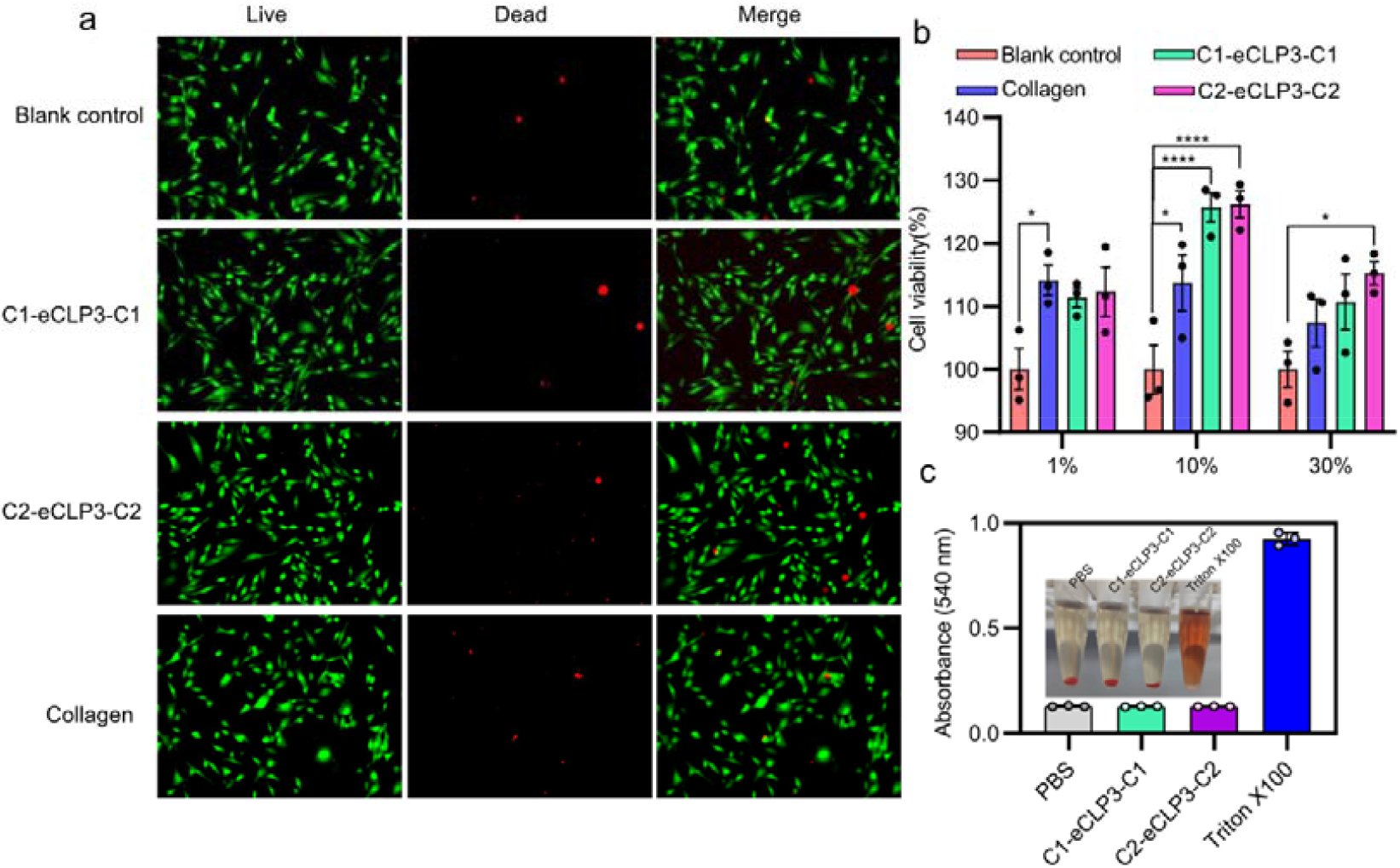
Biocompatibility assessment of hydrogels. a. Representative images of live/dead staining for 3T3 cells treated with C1-eCLP3-C1 and C2-eCLP3-C2 hydrogels. b. Cell viability evaluated using the CCK-8 assay for 3T3 cells treated with C1-eCLP3-C1 and C2-eCLP3-C2 hydrogels. c. Hemolysis analysis of C1-eCLP3-C1 and C2-eCLP3-C2 hydrogels to assess their hemocompatibility. Data are presented as means ± SD. *p < 0.05, **p < 0.01, ***p < 0.001.

### 2.5 Cell Migration Assessment

To further investigate the effects of hydrogels on cell migration, a cell scratch test was conducted. NIH-3T3 cells were scratched using a pipette tip and then treated with 10% hydrogels or native collagen. The cells were imaged at various time points (0-36 hours) after the scratch. The results demonstrated that the 10% hydrogel treatment significantly enhanced the wound recovery compared to the blank control group and the collagen-treated group **(Fig. 5a)**. Interestingly, the C2-eCLP3-C2 hydrogel exhibited greater cell migration ability and cell viability compared to the C1-eCLP3-C1 hydrogel **(Fig. 5b)**. We speculate that this may be attributed to the favorable physical-chemical properties of the C2-eCLP3-C2 hydrogel for cell growth. However, further investigation is required to elucidate the underlying mechanisms involved.

**Figure 5.**
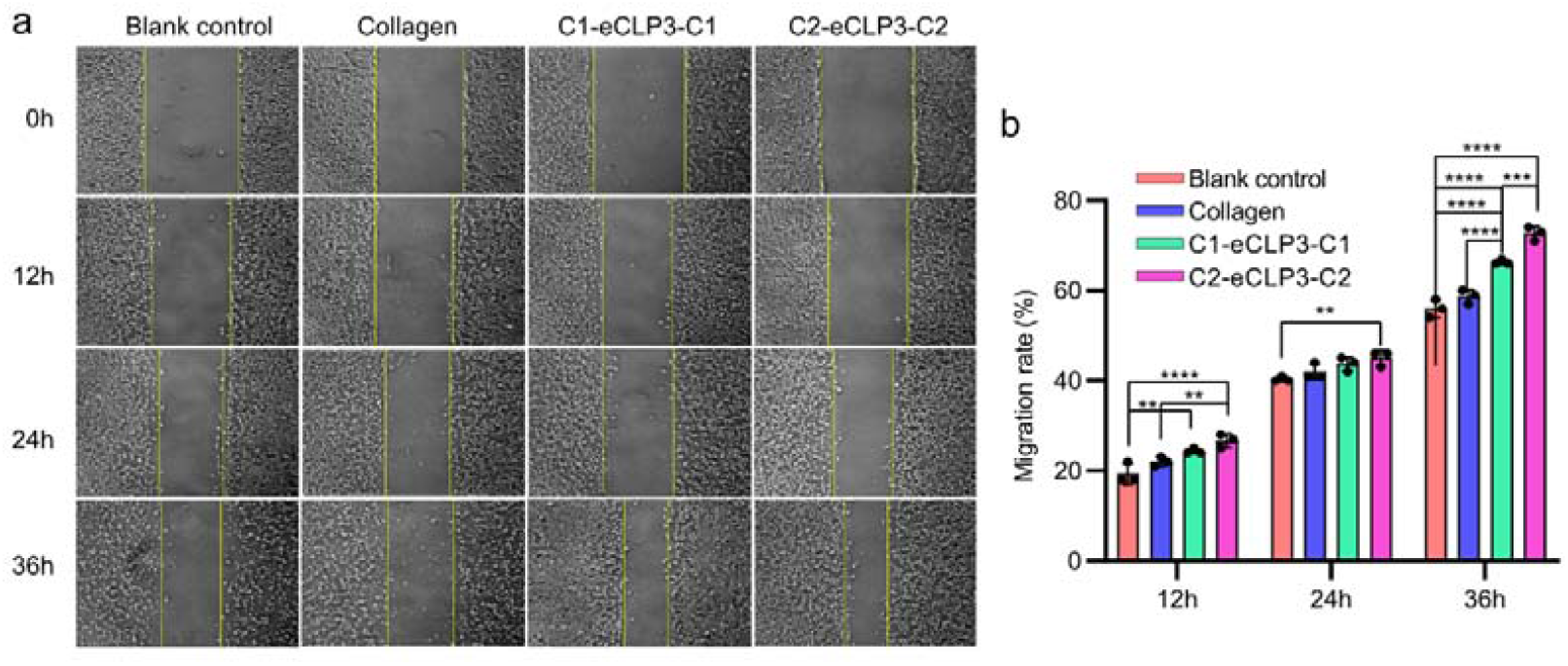
Cell migration assay following hydrogel treatment. a. Representative images showing cell migration at different time points. b. Quantification of cell migration rates. Data are presented as means ± standard deviation (SD). *p < 0.05, **p < 0.01, ***p < 0.001.

### 2.6 Diabetic Wound Healing and Pathological Analysis

The in vitro study provided evidence for the efficacy of C1-eCLP3-C1 and C2-eCLP3-C2 hydrogels in promoting cell growth and viability. To evaluate their therapeutic potential in vivo, a diabetic wound mouse model was established by administering streptozocin (STZ) via intraperitoneal injection for 7 days **(Fig. 6a)**. The induction of diabetes was confirmed by measuring fasting blood glucose levels, which exhibited a significant increase compared to the control group receiving PBS injection, validating the successful establishment of the diabetic mouse model **(Fig. 6b)**. Subsequently, a 1 cm circular wound was created on the dorsum of each mouse. The diabetic wounds were treated with C1-eCLP3-C1 and C2-eCLP3-C2 hydrogels, while PBS was administered to both normal and diabetic mice as control groups. The progression of wound healing was monitored on days 0, 3, 6, 9, and 12 **(Fig. 6c)**. The results demonstrated delayed healing in the PBS-treated diabetic wounds, whereas normal mice exhibited normal wound healing in response to PBS treatment. Importantly, the application of C1-eCLP3-C1 and C2-eCLP3-C2 hydrogels significantly accelerated the rate of healing in diabetic wounds **(Fig. 6c)**. After 12 days of treatment, the wound healing rate reached 80% in the C1-eCLP3-C1 and C2-eCLP3-C2 hydrogel-treated groups, while remaining below 60% in the two control groups **(Fig. 6d)**. At the end of the 12-day monitoring period, all mice were euthanized, and the skin tissue at the wound site was collected for pathological analysis. Hematoxylin-eosin (H&E) and Masson’s trichrome staining were performed to investigate the effects of the hydrogels on wound repair. The hydrogel-treated groups exhibited dense granulation tissues, as well as the formation of newly regenerated epidermis and dermis, while the PBS-treated diabetic mice showed incomplete tissue regeneration **(Supplementary Fig. 2)**. These findings underscore the significant therapeutic potential of C1-eCLP3-C1 and C2-eCLP3-C2 hydrogels in enhancing the healing process of diabetic wounds.

**Figure 6.**
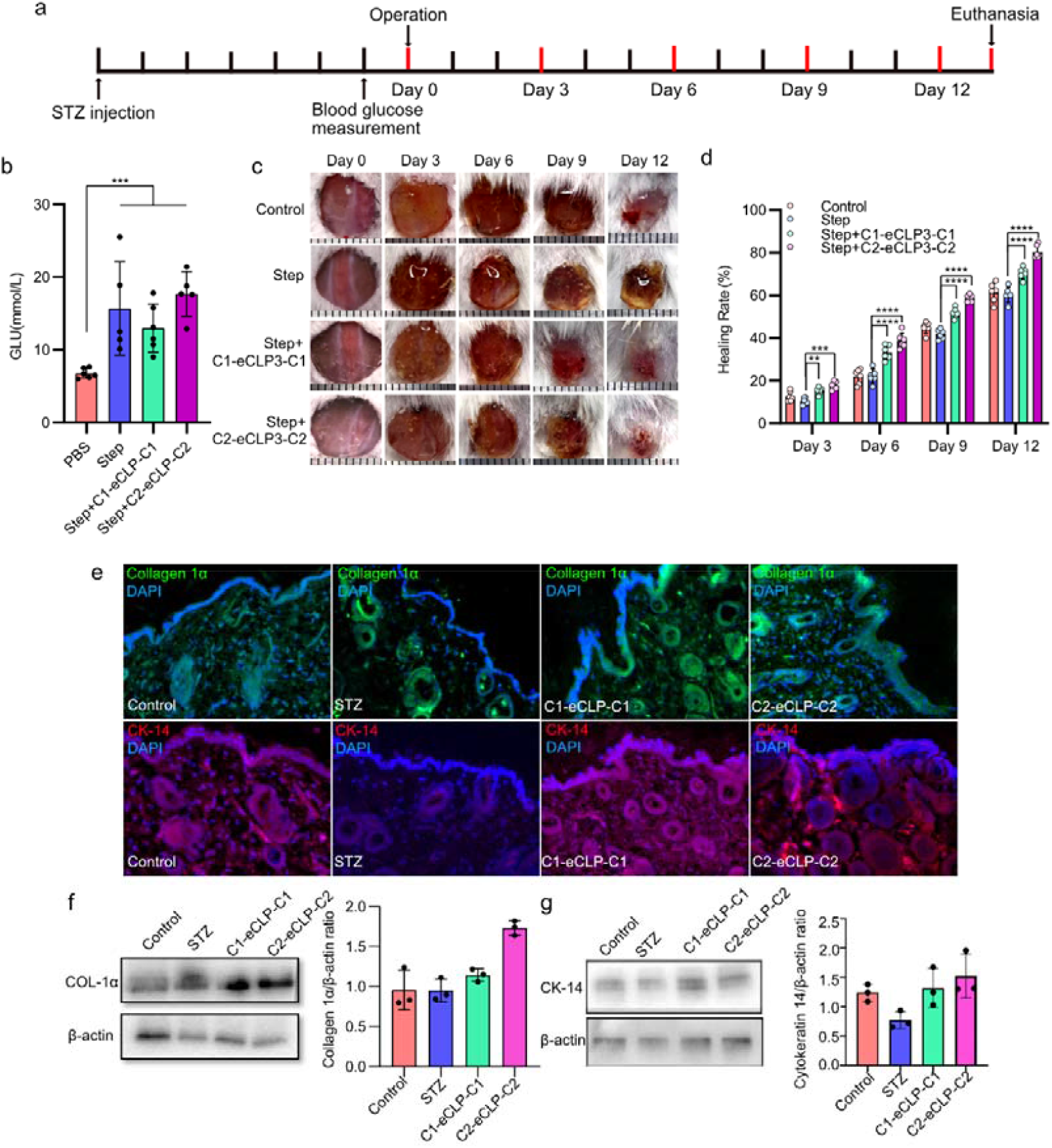
Effects of the hybrid hydrogel on the promotion of diabetic wound healing. a. Experimental timeline for the in vivo study. b. Blood glucose levels in mice after streptozocin (STZ) treatment. c. Representative photographs of wounds in different treatment groups. d. Quantitative analysis of wound area (n ≥ 6). e. Representative immunohistochemistry staining images for COL-1α and CK-14. f. Western blot analysis of COL-1α expression. g. Western blot analysis of CK-14 expression. Data are presented as means ± standard deviation (SD). n = 3, *p < 0.05, **p < 0.01, ***p < 0.001.

To further investigate the wound healing properties of the hydrogels, we conducted immunofluorescence and Western blot analyses to assess the expression of collagen-1α (COL-1α) and cytokeratin-14 (CK-14) at the wound site[8]. Our findings demonstrated a significant upregulation of collagen-1α and CK-14 expression in the wound site upon treatment with C1-eCLP3-C1 and C2-eCLP3-C2 hydrogels, while diabetic wounds exhibited notably lower expression levels of these proteins **(Fig. 6e, f and g)**. These results were further validated by Western blot analysis, which revealed an overall increase in collagen-1α and CK-14 expression in response to C1-eCLP3-C1 and C2-eCLP3-C2 hydrogel treatment **(Fig. 6e, f and g)**. These observations highlight the remarkable ability of C1-eCLP3-C1 and C2-eCLP3-C2 hydrogels to promote epithelial tissue regeneration in diabetic mice, thereby facilitating the wound healing process.

Due to the established roles of cysteine in antioxidant defense mechanisms [31] and electron transfer reactions [32], as well as the reported redox-responsive properties of the hydrogels investigated in this study, we evaluated the levels of superoxide dismutase (SOD) and malondialdehyde (MDA) in the wound site **(Supplementary Fig.3)**. Diabetic wounds have been associated with elevated reactive oxygen species (ROS) production compared to normal wounds [33, 34]. Consequently, in the streptozocin (STZ) treated group, lower SOD levels and higher MDA levels were observed compared to the blank control group. Remarkably, the application of C1-eCLP3-C1 and C2-eCLP3-C2 hydrogels significantly elevated SOD levels and decreased MDA levels at the wound site **(Supplementary Fig.3)**. These findings provide evidence that the C1-eCLP3-C1 and C2-eCLP3-C2 hydrogels possess the ability to reduce excessive ROS production and mitigate oxidative stress in vivo, which is consistent with their observed enhanced wound healing properties.

## 3. Discussion

In this study, we employed genetic encoding techniques to develop two collagen-like proteins based on Scl-2 from *Streptococcus pyogenes* [26]. These proteins were strategically modified with one or two pairs of cysteines, RGD peptides[21], and CPPC peptides[25], resulting in the synthesis of C1-eCLP3-C1 and C2-eCLP3-C2, respectively. Notably, both C1-eCLP3-C1 and C2-eCLP3-C2 proteins exhibited remarkable gelation properties under oxidative conditions or spontaneous gelation after 48 hours of incubation at 4□. The collagen-like protein-based hydrogel derived from these constructs demonstrated a unique gel-sol transition behavior in response to changes in the surrounding environment **(Fig.3)**. This innovative approach holds great promise for the development of advanced biomaterials with tunable properties for various biomedical applications.

The crosslinking strategy is a crucial determinant of the chemical, physical, and biological properties of collagen-like protein-based hydrogels[19, 21, 25]. Various crosslinking methods have been reported for collagen-like protein crosslinking and hydrogel preparation, such as photo-crosslinking in PEG networks[24], EDC-NHS[21], photo-crosslinking[35] and supramolecular approach[19]. However, these methods often involve the use of specific chemicals or modifications that can result in protein denaturation, reduction in protein properties, and limited responsiveness to external stimuli. In contrast, the paired cysteine crosslinking methods we employed in this study are genetically encodable and offer a spontaneous crosslinking approach with redox-responsive properties. This approach minimizes interference with the protein properties and allows for the preservation of their inherent characteristics.

Based on the sulfhydryl of cysteines crosslinking approach, previous studies have reported the use of cysteine-modified elastin with either photo-initiator-induced crosslinking [15] or spontaneous crosslinking[20]. In comparison to these studies, our approach involves the introduction of cysteines into the sequences of collagen-like proteins, resulting in hydrogels with distinct properties compared to elastin-based hydrogels[20]. Furthermore, we assessed the diabetic wound repair ability of collagen-like protein-based hydrogels, which provides valuable insights into the biological effects of collagen-like proteins and their hydrogel formulations.

Based on the findings from both in vitro and in vivo experiments, C1-eCLP3-C1 and C2-eCLP3-C2 exhibited remarkable biocompatibility and demonstrated potential in enhancing wound repair. These positive outcomes can be primarily attributed to the inherent properties of the collagen-like protein itself. The inclusion of the RGD peptide and the protein scaffold within the hydrogel creates an advantageous environment that supports cell growth and migration, eliminating the need for exogenous growth factors or cytokines. To further enhance the reparative effects, it is possible to introduce suitable growth factors or cytokines into the hydrogel by genetically encoding. This is facilitated by the crosslinking strategy employed in our study, which enables protein gelation under mild conditions without the requirement for additional chemical agents. By genetically encoding specific factors into the collagen-like protein-based hydrogel, it becomes feasible to harness the potential of growth factors or cytokines to augment the wound healing process.

Oxidative stress is a detrimental factor associated with diabetes and contributes to the development of chronic wounds in diabetic patients. To address this issue, hydrogels with antioxidative stress capabilities [36] or ROS-scavenging[34] have been developed for the repair of diabetic chronic wounds. Typically, these hydrogels require the addition of exogenous antioxidative chemical compounds to enhance their efficacy. However, in our study, we observed antioxidative effects in C1-eCLP3-C1 and C2-eCLP3-C2 hydrogels without the need for external antioxidative compounds. This can be attributed to the presence of cysteines in the hydrogels, which creates an optimal environment for the stabilization of the sulfhydryl group, as supported by previous studies[31, 32]. The thiolate moiety formed by the cysteines contributes to the observed antioxidative effects[32]. By harnessing the intrinsic properties and design of the reported proteins, our hydrogels exhibit inherent antioxidative capabilities, eliminating the requirement for the addition of external chemical compounds.

Surprisingly, our in vivo experiments revealed that the C2-eCLP3-C2 hydrogel exhibited superior wound repair effects compared to the C1-eCLP3-C1 hydrogel. We observed a significantly higher expression of COL-1α, a key collagen protein, in the diabetic wound site treated with the C2-eCLP3-C2 hydrogel compared to the C1-eCLP3-C1 hydrogel. This difference in COL-1α expression may account for the enhanced wound repair effects observed with the C2-eCLP3-C2 hydrogel. Furthermore, we speculate that the physical mechanisms of the C2-eCLP3-C2 hydrogel are better suited for wound repair compared to the C1-eCLP3-C1 hydrogel, which may contribute to its superior performance. However, the specific underlying mechanisms responsible for these differences require further investigation to provide a comprehensive understanding. In summary, our findings demonstrate the unexpected and superior wound repair effects of the C2-eCLP3-C2 hydrogel compared to the C1-eCLP3-C1 hydrogel in vivo. The elevated expression of COL-1α and the potentially more suitable physical mechanisms of the C2-eCLP3-C2 hydrogel suggest promising avenues for further research to uncover the detailed mechanisms driving these observed differences.

## 4. Materials and Methods

### 4.1 Chemical agents

Scl2 plasmid and primers were synthesized by Sangon Biotech Co.,Ltd and re-constructed by ourself. All used ligases, endonuclease and polymerase was purchased from Thermo Scientific. The antibody of Collagen-1α (CAS: Ab270993) and Cytokeratin-14 (CAS: Ab119695) and the Goat Anti-Mouse IgG H&L (Alexa Fluor® 488) (CAS: Ab150113) were purchased from Abcam. CCK-8 and Live/Dead staining kit was supplied by Beijing Solarbio Science & Technology Co.,Ltd.

### 4.2 Synthesis of plasmids

The Scl2 plasmid was synthesized by Sangon Biotech Co., Ltd. Subsequently, the construction of C1-eCLP3-C1 and C2-eCLP3-C2 sequences was achieved by introducing three repeated collagen-like domains, CGG, RGD, and CPPC sequences were introduced into the plasmid by overlap PCR to construct C1-eCLP3-C1 and C2-eCLP3-C2 sequences, respectively. The protein sequences are showed in supplementary information. pET-28a (+) vector was obtained as a gift from Hermann Gaub (Addgene plasmid # 90473; http://n2t.net/addgene:90473; RRID: Addgene_90473). The PCR production and pET-28a (+) vector were then digested by BamHI and HindIII restriction enzymes of Thermo Scientific. The insertions and vector then ligated by T4 ligase of Thermo Scientific.

### 4.3 Recombinant Protein Expression in E. coli

The plasmids containing C1-eCLP3-C1 and C2-eCLP3-C2 sequences were transformed into the BL21(DE3) *E. coli* strain. After verification and sequencing, the plasmids were expressed in BL21(DE3) *E. coli* cells. Protein expression was conducted in shake flasks using 6 L of Luria-Bertani liquid medium (LB medium) supplemented with 50 μg/mL of kanamycin at 37 °C. Once the optical density at 600 nm (OD_600_) reached 0.8, the cells were induced with β-D-Thio-galactoside (IPTG) at a final concentration of 0.3 mM and incubated overnight at 16 °C. After a 32-hour incubation period, the cells were harvested by centrifugation at 8000g for 15 minutes at 4 °C.

### 4.4 Recombinant Protein Purification

After centrifugation, the cell pellet was resuspended in a lysis buffer (50 mM Tris-HCl, pH 8.0, 300 mM NaCl, 5% glycerol, v/v). The resuspended cells were then subjected to ultrasonication to disrupt the cell membranes, followed by centrifugation at 12,000 rpm for 60 minutes at 4°C. The resulting pellet was resuspended in a binding buffer and filtered through a 0.45 μm filter to remove any remaining debris. The protein of interest was subsequently purified using a Ni^2+^ affinity column, employing a gradient elution with 300 mM imidazole in a buffer containing 50 mM Tris-HCl (pH 8.0), and 300 mM NaCl. The eluted protein was dialyzed against buffer Q (20 mM Tris-HCl, pH 8.0, 50 mM NaCl, 1 mM EDTA, 1 mM Dithiothreitol (DTT), and 5% glycerol, v/v) and loaded onto a HiTrap Capto Q column (GE Healthcare). The protein was further purified using a gradient of NaCl (0-1000 mM) in buffer Q. The purity of the protein was verified by sodium dodecyl sulfate-polyacrylamide gel electrophoresis (SDS-PAGE) followed by Coomassie Brilliant Blue G-250 staining. Additionally, the purified protein was confirmed using Western Blot analysis with an anti-Histag antibody.

### 4.5 Circular Dichroism (CD)

To analyze the protein’s secondary structure and denaturation temperature, we employed circular dichroism (CD) spectroscopy. A Bio-Logic MOS-500 circular dichroism spectropolarimeter was used to measure CD spectra of 10 μM aqueous protein solutions in acidic, neutral, and basic pH buffers. The CD signals were recorded across a wavelength range of 185 to 260 nm at a constant temperature of 25 °C. To ensure accuracy, we collected and averaged ten scans for each pH condition. Additionally, we performed wavelength scans on protein samples at pH 7 in sodium phosphate buffer, covering the range of 185 to 260 nm.

### 4.6 Hydrogel Fabrication and redox-response

The purified proteins were concentrated to a concentration of 20 mg/mL using a 10 kDa Amicon® tube (Millipore, UFC9010), and the buffer was exchanged to PBS. The concentrated proteins were then subjected to incubation at 4 °C for at least 24 hours to facilitate gel formation. To investigate the redox-responsive capability of the protein hydrogels, For the redox-response examination, the hydrogels were initially treated with 100 μM 1,4-Dithiothreitol (DTT), resulting in the hydrogel transitioning to a solution within 10 minutes. Subsequently, a 200 μM H_2_O_2_ solution was added to the protein solution, leading to gel formation after 6 hours of incubation.

### 4.7 Hydrogel Characterization

The protein hydrogel samples were subjected to an overnight incubation at -20 °C, followed by a 24-hour lyophilization process. The resulting lyophilized hydrogel samples were examined for surface topography using a Hitachi S-4800 scanning electron microscope (SEM). To analyze the chemical composition, a pellet of the lyophilized hydrogel, diluted to 1% weight, was prepared and subjected to analysis using a Thermerfeld NICOLET 6700 spectrometer to obtain FT-IR spectra. The spectra were recorded in the range of 4000-400 cm^-1^ with a resolution of 0.1 cm^-1^. Furthermore, the rheological properties of the hydrogel were assessed using a DHR-1 rheometer (TA Company, USA). Various techniques including frequency sweep, time sweep, and strain sweep were employed to evaluate the rheological behavior of the hydrogel.

### 4.8 Cell Viability and Migration

To evaluate cell viability, the hydrogels were subjected to 30 minutes of UV sterilization. NIH-3T3 cells were cultured in DMEM (Gibco, Catalog no. 11966025) medium supplemented with 1% penicillin/streptomycin (Gibco, Catalog no. 2441834) and 10% fetal bovine serum (Gibco, Catalog no. 10100147) at 37 °C in a 5% CO2 atmosphere. After 24 hours, the protein hydrogels were added to the cell culture medium at concentrations of 10% or 30% (v/v) and co-incubated with the cells for 24 hours. For control groups, the medium was supplemented with Type I collagen at the same concentration as the hydrogels. To assess cytotoxicity, live/dead staining was performed on the cells treated with the hydrogels. Live cells were stained with a green fluorescent dye, while dead cells were stained with a red fluorescent dye. This staining allowed for the visualization and quantification of cell viability. In addition, cell viability of NIH-3T3 cells co-cultured with different hydrogels was assessed using a Cell Counting Kit-8 (CCK-8) assay. Specifically, 10,000 cells were seeded in each well of a 96-well plate. After co-culturing for 12 hours, the culture medium was replaced with fresh medium, and the CCK-8 reagent was added to each well. Following a 1-hour incubation, the optical density (OD) was measured at 450 nm, and the results were normalized to the medium control. All experiments were performed in triplicate, and the hydrogels were evaluated for cytotoxicity and their impact on cell viability. To assess cell migration following hydrogel administration, cells were cultured in 24-well plates and subjected to scratch assays using a 1.5 mL pipette tip. Subsequently, hydrogels were added to the culture and co-incubated with the cells. Time-lapse images of the cells were captured at 0, 12, 24, and 36 hours. The obtained images were subsequently analyzed using Fiji software to quantitatively evaluate cell migration.

### 4.9 Hemolysis Test

The hemolysis test was conducted following a previously described method[16]. In brief, 1 mL of whole blood was added to a 1.5 mL Eppendorf tube containing 150 μL of citric acid. The solution was then centrifuged at 3500 rpm for 10 minutes at 4 °C to isolate the red blood cells (RBCs). The RBCs were subjected to three rounds of centrifugation, and then diluted with 20 mL of phosphate buffered saline (PBS) for subsequent use. For the hemolysis test, 200 μL of the RBC suspension was added to 800 μL of the hydrogel suspension. All suspensions were incubated in a rocking shaker at 37 °C for 3 hours, followed by centrifugation at 10,016 × g for 3 minutes. The absorbance of the released hemoglobin in the suspensions was measured at 540 nm using a UV–vis spectrometer. Each experiment was repeated three times to ensure accuracy and reproducibility.

### 4.10 Mouse diabetic wound model

Male Balb/c mice weighing 20-25g were obtained from Charles River and allowed to acclimatize to the laboratory environment for 7 days prior to the experiment. Diabetes was induced by one time for one day intraperitoneal injection of 110 mg kg^−1^ streptozotocin (STZ, Sigma Aldrich) dissolved in citrate buffer (pH 4.5)[36]. After 7 days of STZ treatment, blood glucose levels in the mice were measured. Mice with consistently elevated glucose levels above 10.0 mmol L−1 were considered diabetic. Once the diabetic mouse model was established, the mice were anesthetized using isoflurane (RWD, R510-22) administered via a respiratory anesthesia system (RWD, R510IP). A circular wound with a diameter of 1 cm was created on each mouse. Subsequently, C1-eCLP3-C1 and C2-eCLP3-C2 hydrogels, which had undergone UV sterilization, were applied to the wounds. On days 0, 3, 6, 9, and 12 after treatment, the mice were anesthetized and imaged. The wound areas were analyzed using Fiji software, and the wound healing rate was calculated using the following formula: Wound healing rate (%) = (S0 - Sn) / S0 × 100, where S0 and Sn represent the wound area on day 0 and day n after treatment[37]. All animal experiments were conducted in accordance with the guidelines and regulations approved by the Inner Mongolia University Ethics Committee.

### 4.11 Histology and immunohistochemistry

After 12 days, the mice in each group were euthanized with CO_2_ and the wound skin area was obtained. The wound tissues were fixed in 4% paraformaldehyde for 24 h and dehydrated in sucrose gradient solution. The tissues were then embedded in paraffin and subjected to routine haematoxylin-eosin (H&E) staining and Masson’s trichrome staining. The stained sections were observed using an optical microscope (Leica, DMi8 S). For immunohistochemistry, the methods previously reported were followed [38]. Heat-mediated antigen retrieval was performed for 1 hour at 95 °C, and the sections were then incubated with primary antibodies targeting collagen-1α and cytokeratin-14 at a temperature of 4 °C. Subsequently, the sections were treated with a secondary antibody (anti-rabbit/anti-mouse Envision) for 30 minutes. Finally, fluorescence microscopy (Leica, DMi8 S) was utilized to visualize and analyze the immune-stained sections.

### 4.12 Statistical analysis

The wound site was lysed, and total protein was extracted. The protein concentration was determined using a BCA protein assay kit. Subsequently, 30 µg of protein samples were separated by 10% SDS-PAGE and transferred onto nitrocellulose membranes. Primary antibodies against COL-1α and CK-14 (dilution 1:500, Abcam) were incubated with the membranes overnight at 4 °C, while β-actin served as the internal reference. The membranes were then incubated with a secondary antibody, anti-rabbit IgG horseradish peroxidase-conjugated (dilution 1:5000). Protein bands were visualized using an Enhanced Pico Light Chemiluminescence Kit. Densitometric analysis of the protein bands was performed using Image J software.

The antioxidant effect of the hydrogels was assessed using SOD assay kit and MDA assay kit. Firstly, the protein solution, which was prepared in the same manner as for the Western Blot experiment, was quantified using a BCA protein assay kit. Subsequently, the activity of superoxide dismutase (SOD) and the level of malondialdehyde (MDA) were analyzed according to the instructions provided with the assay kits.

### 4.13 Statistical analysis

In this study, the data are presented as mean ± standard deviation. Statistical analysis was conducted using GraphPad Prism 8 software, utilizing one-way analysis of variance (ANOVA). A p-value of less than 0.05 was considered statistically significant, indicating a significant difference between the groups. Image analysis was performed using Fiji software, enabling accurate and reliable measurements and quantification of the obtained images.

## 5. Acknowledgement

This project was supported by a grant from Inner Mongolia Natural Science Foundation (2022QN03014); Inner Mongolia Science and Technology Project (2022ZY0050); Inner Mongolia Youth Science and Technology Talent Support Project (NUYT23092); Science and Technology Leading Talent Team in Inner Mongolia Autonomous Region (2022LJRC0009). S.J, J.W and X.W contributed equally to this work. All authors have approved the final version of this manuscript. Ethical approval for the in vivo diabetic wound healing experiments was granted by the Institutional Animal Care and Use Committee of Inner Mongolia University.

## 8. Data Availability Statement

The data that support the findings of this study are available from the corresponding author upon reasonable request.

