## Supplementary information for "Genetically Encodable *in situ* Gelation Redox-Responsive Collagen-Like Protein Hydrogel for Accelerating Diabetic Wound Healing"

### Supplementary figure 1: Amino acid sequence of proteins

**C1-eCLP3-C1**

MCGGADEQEEKAKVRTELIQELAQGLGGIEKKNFPTLGDEDLDHTYMTKLLTYLQEREQAENSWRKRLLKGIQDHALDGGPCPPCGPKGEQGPQGLPGKDGEAGAQGPAGPMGPAGEQGEKGEPGTQGAKEDRGETGPKGPKGERGEAGPAGKDGEPGPVGPAGPKGEQGPQGLPGKDGEAGAQGPAGPMGPAGEQGEKGEPGTQGAKEDRGETGPKGPKGERGEAGPAGKDGEPGPVGPAGPKGEQGPQGLPGKDGEAGAQGPAGPMGPAGEQGEKGEPGTQGAKEDRGETGPKGPKGERGEAGPAGKDGEPGPVGPAGGPCPPCRGDGGC*

**C2-eCLP3-C2**

MCGGCGGADEQEEKAKVRTELIQELAQGLGGIEKKNFPTLGDEDLDHTYMTKLLTYLQEREQAENSWRKRLLKGIQDHALDGGPCPPCGPKGEQGPQGLPGKDGEAGAQGPAGPMGPAGEQGEKGEPGTQGAKEDRGETGPKGPKGERGEAGPAGKDGEPGPVGPAGPKGEQGPQGLPGKDGEAGAQGPAGPMGPAGEQGEKGEPGTQGAKEDRGETGPKGPKGERGEAGPAGKDGEPGPVGPAGPKGEQGPQGLPGKDGEAGAQGPAGPMGPAGEQGEKGEPGTQGAKEDRGETGPKGPKGERGEAGPAGKDGEPGPVGPAGGPCPPCRGDGGCGGC*

Amino acid sequence of C1-eCLP3-C1. CGG; V-domain; CPPC; Collagen-like domain-1; Collagen-like domain-2; Collagen-like domain-3; RGD.

### Supplementary Table 1: Proteomics of eCLP3 template protein

| Protein description | Score | Coverage | # Peptides | # PSMs | # AAs | MW [kDa] | calc. pI |
| --- | --- | --- | --- | --- | --- | --- | --- |
| eCLP3 | 6442.00 | 73.00 | 20 | 199 | 363.00 | 36.40 | 5.90 |
| Sequence | PSMs | Modifications | IonScore | Charge | MH+[Da] | ΔM[ppm] | RT [min] |
| DGEPGPVGPAGPK | 40 |  | 53 | 2 | 1177.58 | 0.51 | 25.23 |
| DGEPGPVGPAGPKGEQGPQGLPGK | 1 |  | 46 | 2 | 2226.11 | -0.17 | 12.31 |
| EQAENSWR | 1 |  | 39 | 2 | 1019.45 | 0.23 | 8.31 |
| GEAGPAGKDGEPGPVGPAGPK | 11 |  | 98 | 2 | 1844.91 | 2.66 | 9.69 |
| GEPGTQGAK | 1 |  | 31 | 2 | 844.42 | -0.99 | 5.66 |
| GEPGTQGAKEDRGETGPK | 2 |  | 55 | 2 | 1813.87 | 1 | 6.69 |
| GEQGPQGLPGK | 17 |  | 37 | 2 | 1067.55 | 2.08 | 9.34 |
| GEQGPQGLPGKDGEAGAQGPAGPMGPAGEQGEK | 2 |  | 81 | 2 | 3059.41 | 0.83 | 12.32 |
| GEQGPQGLPGKDGEAGAQGPAGPMGPAGEQGEK | 2 | 1xOxidation[M24] | 36 | 3 | 3075.41 | 1.18 | 10.78 |
| GEQGPQGLPGKDGEAGAQGPAGPMGPAGEQGEKGEPGTQG | 1 | 1xOxidation[M24] | 32 | 3 | 3900.81 | 1.77 | 10.53 |
| GIQDHALDGGPCPPCR | 1 | 2xCarbamidomethyl[C12;C15] | 24 | 3 | 1749.78 | -0.26 | 11.38 |
| GPKGEQGPQGLPGK | 2 |  | 67 | 2 | 1349.72 | 0.65 | 8.28 |
| KNFPTLGDEDLDHTYMTK | 1 |  | 61 | 3 | 2124.99 | 2.38 | 15.02 |
| KNFPTLGDEDLDHTYMTK | 2 | 1xOxidation[M16] | 56 | 2 | 2140.99 | 1.29 | 13.24 |
| LLTYLQER | 43 |  | 59 | 2 | 1035.58 | -0.02 | 14.76 |
| LLTYLQEREQAENSWR | 1 |  | 26 | 3 | 2036.02 | -0.52 | 15.57 |
| NFPTLGDEDLDHTYMTK | 5 |  | 131 | 2 | 1996.90 | 5.01 | 16.96 |
| NFPTLGDEDLDHTYMTK | 11 | 1xOxidation[M15] | 91 | 2 | 2012.89 | 0.46 | 16.45 |
| TELIQELAQGLGGIEK | 36 |  | 93 | 2 | 1698.93 | 0.75 | 24.03 |
| TELIQELAQGLGGIEKK | 2 |  | 73 | 2 | 1827.02 | -0.5 | 22.03 |
| VRTELIQELAQGLGGIEK | 1 |  | 95 | 2 | 1954.10 | -1.39 | 21.90 |

### Supplementary figure 2: Amino acid sequence of proteins

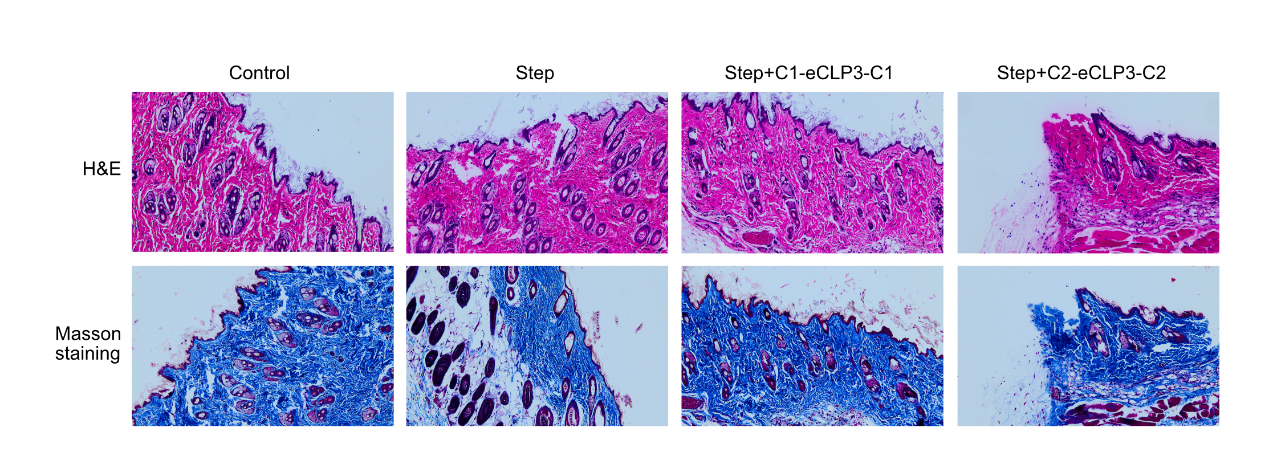

Supplementary figure2. Hematoxylin-eosin (H&E) and Masson's trichrome staining were performed to investigate the effects of the hydrogels on wound repair.

### Supplementary figure 3: Amino acid sequence of proteins

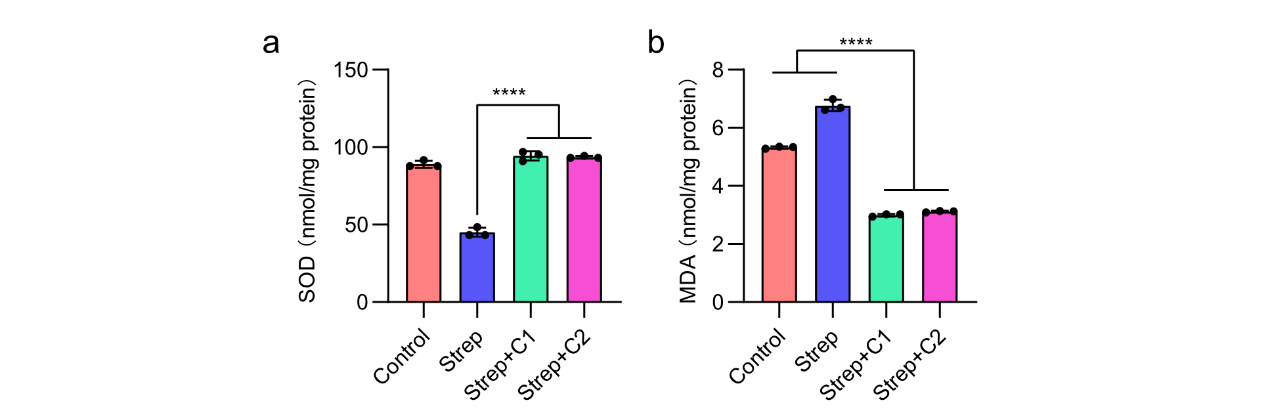

**Supplementary figure 3**. Superoxide Dismutase (SOD) and Malondialdehyde (MDA) levels at the diabetic wound site. a. SOD levels at the wound site. b. MDA levels at the wound site.
